# Influence of molecular representation and charge on protein-ligand structural predictions by popular co-folding methods

**DOI:** 10.64898/2026.02.18.706547

**Authors:** Alisa Bugrova, Philipp Orekhov, Ivan Gushchin

## Abstract

Recently developed deep learning-based tools can effectively generate structural models of complexes of proteins and non-proteinaceous compounds. While some of their predictive capabilities are truly exciting, others remain to be thoroughly tested. Here, we probe whether the ligand input format (Chemical Component Dictionary, CCD, or Simplified Molecular Input Line Entry System, SMILES) and charge (which depends on protonation) will affect the results of the predictions by four popular algorithms: AlphaFold 3, Boltz-2, Chai-1, and Protenix-v1. We chose methylamine and acetic acid as two of the simplest titratable chemicals that are omnipresent in proteins as amino and carboxy moieties, and are consequently ubiquitous in the Protein Data Bank models that are most commonly used for training. Unexpectedly, we found that for both molecules, in many cases the input format affected the prediction results, and did it much stronger compared to protonation, whereas changes in the formally specified charge of the molecules did not lead to changes in binding expected from experiments. We conclude that (i) ensuring identical results irrespective of input formats and (ii) inclusion of protonation-related steps into training and prediction pipelines are the two available paths for improvement of protein-ligand structure prediction algorithms.

## Introduction

Protein structure prediction using deep learning revolutionized the field of structural biology by allowing mass generation of realistic models and expanding our understanding of patterns that govern interactions between residues and proteins [1]. New tools, such as AlphaFold 3 [2], also called co-folding methods, are trained to predict models of assemblies of biomolecules of different types, including proteins-ligand complexes. These capabilities are promising not only for better understanding of basic biology, but also for biotechnology and drug development [2].

One of the main limitations of AlphaFold 3 is that despite being the first state-of-the-art model capable of handling small molecules, it is not universally available and has a restrictive license. Several other models address these issues, such as Boltz, Chai and Protenix. Chai-1 is an openly accessible model that follows Alphafold 3-inspired architecture and training strategy [3]. Protenix is also one such model, claiming in its most recent implementation (Protenix-v1) to reach and exceed AlphaFold 3 level [4, 5]. Finally, Boltz-2 additionally introduced physics-based potentials and a separate model for affinity prediction, aimed at facilitating development of therapeutics [6].

Reports that introduce the models, as well as general benchmarks, report relative success of protein-ligand complex prediction with AlphaFold 3 and its analogs [2–9]. However, systematic errors are also observed. For example, distributions of amino acid bond length and angles differ from those observed in experiments [10]; ligands and generated complexes may contain stereochemical errors [11]. Structures of protein-ligand complexes appear to be memorised to some extent: success rate for molecules dissimilar to those found in the training set is statistically lower [12, 13]. The ligands may be positioned similarly by the algorithms despite drastic alterations in the receptor (binding site removal, overpacking, charge inversion) [14]. Ligands with opposing implied charges are also positioned similarly, which contradicts basic physico-chemical principles [14].

Here, we decided to test whether the results of predictions by popular widely available cofolding tools, AlphaFold 3, Boltz-2, Chai-1 and Protenix-v1, will be sensitive to explicitly specified ligand charges. We chose methylamine and acetic acid as two of the simplest titratable chemicals that are omnipresent in proteins as amino- and carboxy-moieties, thus ubiquitous in the available training sets and expected to be modeled well. We tested SMILES, a notation that encodes the molecule as a string [15] and CCD, a Chemical Component Dictionary code [16], as the input formats that allow to specify the charge of the molecule. Unexpectedly, we found that in many algorithm-molecule combinations, the input format affected the results of the prediction, whereas charge did not.

## Methods

Inference for all structure prediction models was run on the NVIDIA GeForce RTX 4090 Graphics Processing Unit (GPU) with locally installed models.

AlphaFold 3 [2] v. 3.0.1 was run locally using HMMER 3.4 [17] for Multiple Sequence Alignment (MSA) generation, installed as described in the Dockerfile. MSA generation was first performed for a protein without ligand, and then the MSA without templates was transferred to the input JavaScript Object Notation (JSON) files for complex predictions. Structure prediction was run for 100 different random seed values with 5 diffusion samples generated per seed for each ligand and input format.

Boltz-2, Chai-1 and Protenix-v1 accessed a local Colabfold MSA server [18] configured with GPU but without GPU-server for MSA generation. For all three programs, the MSAs were first generated during structure prediction without ligands and then used for complex prediction. Boltz-2 [6] v. 2.2.1 was run with the default version 2 weights. For BarA, 500 diffusion samples were generated at once for each ligand and input format. For the dopamine receptor, the structure prediction for each ligand and input format was performed in 5 runs comprising 100 diffusion samples each. Chai-1 [3] v. 0.5.2 was run with the default weights 50 times with 10 diffusion samples for each ligand. Given that Chai-1 only accepts SMILES as the input format, CCD input was not tested. For Protenix-v1 [5] the structure prediction was run with the 1.0.0 version of the code and default weights 50 times with seeds from 1 to 50 and 10 samples for each ligand and input format.

To facilitate the predictions and limit the modeled region to ordered ligand-binding domain, we used residues 21-346 from the UniProt ID P21728 sequence for dopamine D1 receptor predictions and residues 31-178 from the Uniprot ID P0AEC5 sequence for BarA predictions [19]. The CCD codes and SMILES strings used for each ligand were as follows: NME and CN for methylamine, 3P8 and C[NH3+] for methylammonium, ACY and CC(=O)O for protonated acetic acid, ACT and CC(=O)[O-] for acetate. The CCD codes were obtained from PDBeChem [20] and the SMILES strings were obtained from the PubChem database [21].

Structure analysis was performed using the Biopython PDB module [22]. All models for a certain protein target were superimposed based on the protein backbone atoms onto the structure of the same target generated in Alphafold 3 with no ligand. After that, the ligand atom coordinates were extracted and analysed. Principal component analysis (PCA) was performed using the Scikit-learn Python module [23]. The Kolmogorov–Smirnov statistics were calculated using the SciPy Python module [24]. Data analysis and visualisation was performed with the Pandas [25] 2.2.2 and Plotly [26] 6.3.0 Python modules. Structure visualisation was performed with PyMOL [27] 2.5.0 Open-Source.

## Results

To assess the performance of the cofolding methods depending on explicitly specified charge of the ligand and on the input format, we have chosen human dopamine D1 receptor (DRD1) for methylamine and methylammonium tests and *E. coli* BarA kinase sensor domain for the acetate and acetic acid tests. DRD1 is an important clinical target as it is central in multiple dopaminergic pathways in the human neural system [28]. Binding of dopamine and chemically related molecules to the receptor involves formation of an ionic bridge between the amino group of the ligand and the carboxylate group of D103 [28]. Accordingly, charged methylammonium ion is expected to be positioned near D103 by the algorithms, whereas neutral methylamine is likely to be bound elsewhere, if at all. BarA is a receptor histidine kinase that participates in gene expression regulation in *E. coli*. Its sensor domain binds protonated forms of acetate and formate that serve as physiological stimuli [29]. Putative ligand binding pocket of BarA is composed of hydrophobic and polar residues L95, T98, and I136 [29]. Accordingly, neutral acetic acid is expected to be positioned near these residues, whereas negatively charged acetate ion should not occupy the pocket.

First, we analyzed the predicted structures of the ligands themselves. Theoretical lengths of the C-N bonds calculated using quantum chemical approaches are 1.45-1.48 Å for methylamine and 1.48-1.51 Å for methylammonium [30–33]. We found that the predicted bond lengths are on average shorter, falling out of the expected ranges (Fig. 1, Table 1). Surprisingly, Protenix-v1 generated bond lengths as short as 0.075 Å, with 65 values for methylammonium out of 1000 analyzed being relatively evenly distributed in the completely unnatural range from 0 to 1 Å. Chai-1 predictions are not sensitive to charge, whereas AlphaFold 3, Boltz-2 and Protenix-v1 produce different length distributions depending on charge; the sign of the difference is correct only in one case (Boltz-2 with input specified using SMILES).

**Table 1.** Means and standard deviations of interatomic distances for various ligand and input format combinations predicted by different tools. O1 and O2 denote oxygen atoms with the smaller and larger index, respectively. Reference ranges are inferred from quantum chemical calculations [30–37].

|  | AlphaFold 3 | Boltz-2 | Chai-1 | Protenix-v1 | Reference |
| --- | --- | --- | --- | --- | --- |
| N-C distance, Å |  |  |  |  |  |
| Methylamine CCD | 1.319±0.015 | 1.260±0.007 |  | 1.261±0.028 | 1.45 - 1.48 |
| Methylamine SMILES | 1.306±0.017 | 1.258±0.005 | 1.433±0.042 | 1.293±0.049 | 1.45 - 1.48 |
| Methylammonium CCD | 1.273±0.023 | 1.263±0.005 |  | 1.205±0.175 | 1.48 - 1.51 |
| Methylammonium SMILES | 1.273±0.023 | 1.281±0.005 | 1.454±0.038 | 1.193±0.183 | 1.48 - 1.51 |
| C-C distance, Å |  |  |  |  |  |
| Acetate CCD | 1.469±0.007 | 1.416±0.027 |  | 1.481±0.010 | 1.52 - 1.534 |
| Acetate SMILES | 1.461±0.004 | 1.453±0.004 | 1.465±0.019 | 1.456±0.009 | 1.52 - 1.534 |
| Acetic Acid CCD | 1.491±0.008 | 1.433±0.024 |  | 1.484±0.010 | 1.495 - 1.502 |
| Acetic Acid SMILES | 1.464±0.004 | 1.448±0.005 | 1.471±0.018 | 1.456±0.009 | 1.495 - 1.502 |
| C-O distance, Å |  |  |  |  |  |
| Acetate CCD<br>C-O<br>C-OXT | 1.224±0.006<br>1.258±0.008 | 1.219±0.011<br>1.209±0.036 |  | 1.212±0.013<br>1.203±0.011 | 1.26 - 1.28<br>1.26 - 1.28 |
| Acetate SMILES<br>C-O1<br>C-O2 | 1.242±0.003<br>1.243±0.004 | 1.212±0.006<br>1.231±0.006 | 1.262±0.018<br>1.254±0.018 | 1.226±0.014<br>1.222±0.008 | 1.26 - 1.28<br>1.26 - 1.28 |
| Acetic Acid CCD<br>C-O<br>C-OXT | 1.237±0.005<br>1.254±0.006 | 1.223±0.010<br>1.203±0.009 |  | 1.212±0.012<br>1.211±0.012 | 1.189 - 1.22<br>1.331 - 1.35 |
| Acetic Acid SMILES<br>C-O1<br>C-O2 | 1.245±0.003<br>1.248±0.003 | 1.220±0.008<br>1.229±0.007 | 1.268±0.018<br>1.258±0.017 | 1.235±0.012<br>1.234±0.012 | double bond<br>1.189 - 1.22<br>single bond<br>1.331 - 1.35 |

**Figure 1.**
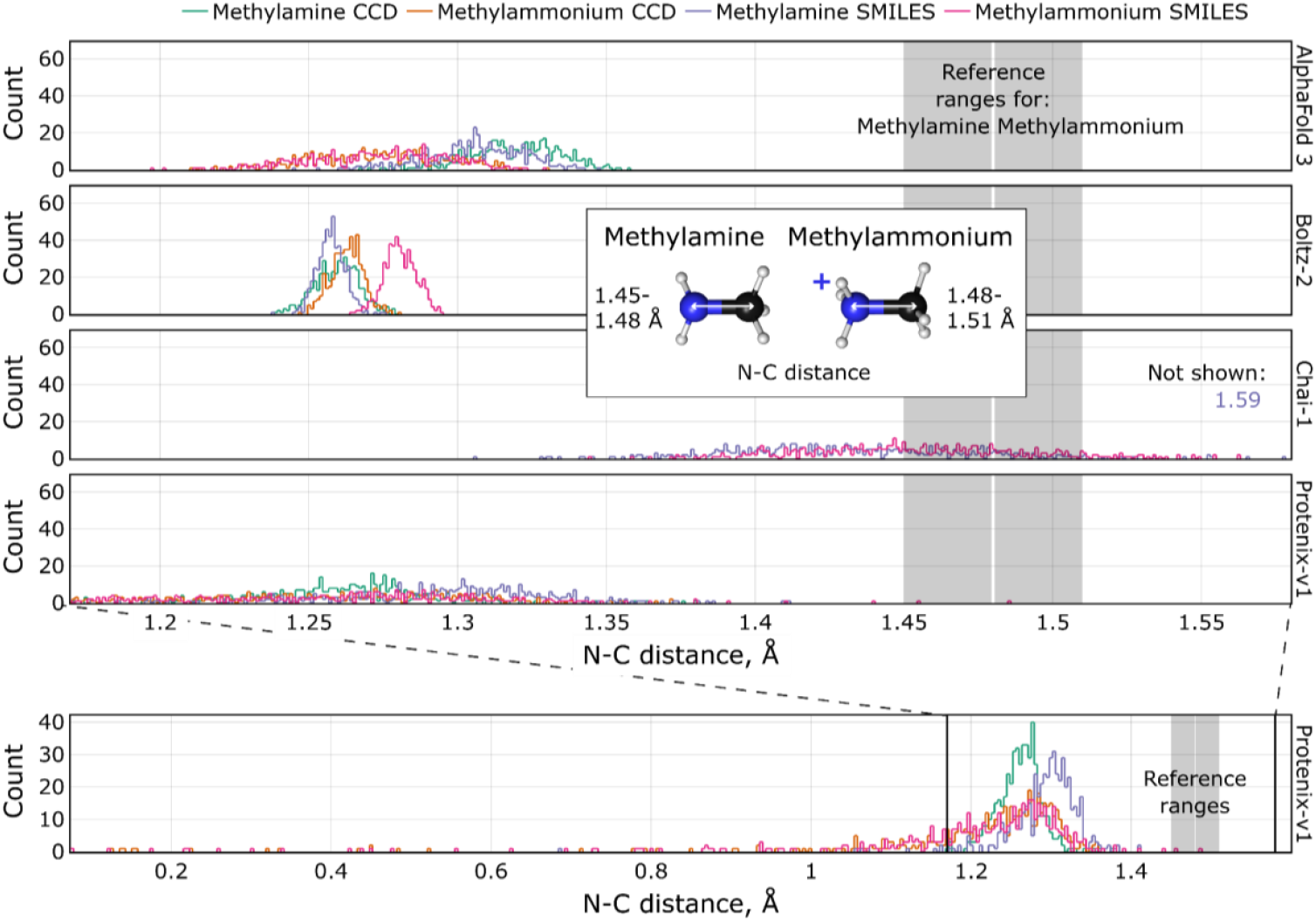
Distributions of predicted distances between the heavy atoms in methylamine and methylammonium. Reference ranges obtained using quantum chemical calculations [30–33] are highlighted in gray.

For acetate, the C-C bond length is expected to lie within 1.52-1.534 Å, whereas C-O bond lengths are expected to be identical and lie within 1.26-1.28 Å [30, 34, 35]. For protonated acetic acid, the C-C bond length is expected to lie within 1.495-1.502 Å, the C-O single bond within 1.331-1.35 Å and the C-O double bond within 1.189-1.22 Å [30, 35–37]. For all of the bonds, we find that Chai-1 predictions display the most variance, and Chai-1 and Protenix-v1 predictions are not sensitive to charge (Fig. 2A, Table 1). Predicted C-C bond lengths are again on average shorter than expected. AlphaFold 3 and Boltz-2 produce different C-C length distributions depending on charge; the sign of the difference is correct only in one case (Boltz-2 with input specified using SMILES). As for the C-O bonds, we note that in CCD notation, the single-bonded oxygen is denoted as OXT and the double-bonded oxygen is denoted as O. In protonated acetic acid, the two C-O distances are expected to be different. We do not find any pattern in predicted bond lengths; somehow the two bond lengths are often more different for acetate rather than for acetic acid (Fig. 2B). Only the Chai-1 predictions for acetate cluster around the expected values. For Protenix-v1, some C-O bond length distributions are noticeably not unimodal (Fig. 2B).

**Figure 2.**
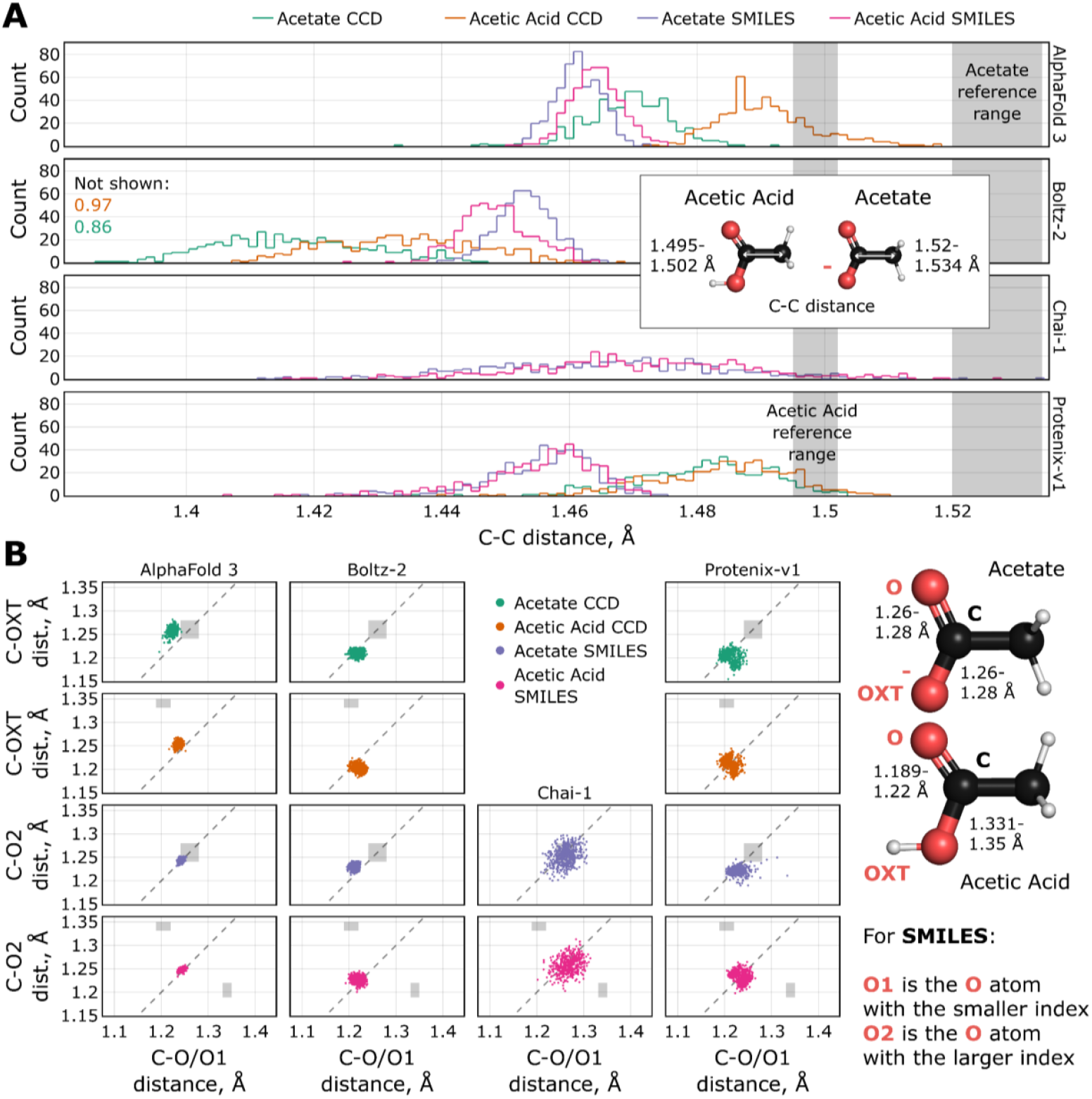
Distributions of predicted distances between the heavy atoms in acetate and acetic acid. A) C-C distance. B) C-O distances. The dashed line corresponds to identical C-O distances within the molecule. In some cases, systemic bias is observed. Reference ranges obtained using quantum chemical calculations [30, 34–37] are highlighted in gray. C-O distances are expected to be similar in acetate and systematically different in acetic acid.

Next, we analyzed the predicted ligand positions relative to the protein. For DRD1 complexed with methylammonium and methylamine, the ligands are mostly positioned close to the expected amino group location observed in experiments (Fig. 3), whereas in Chai-1 and Protenix-v1 predictions the positions are increasingly diverse, with some ligands placed far from the expected position. In order to quantify the differences, we analyzed the predicted positions using PCA [38], and compared the distributions along PC1 using the Kolmogorov-Smirnov statistic, equal to 0 for identical distributions and to 1 for non-overlapping distributions [39]. AlphaFold 3 and Protenix-v1 predictions for methylammonium do not depend on the input format, whereas distributions for methylamine are different for CCD and SMILES; Boltz-2 predictions are extremely sensitive to the input format. At the same time, position distributions for all methods except Boltz-2 are mostly independent of charge, whereas for Boltz-2 they are somewhat dependent (Fig. 3).

**Figure 3.**
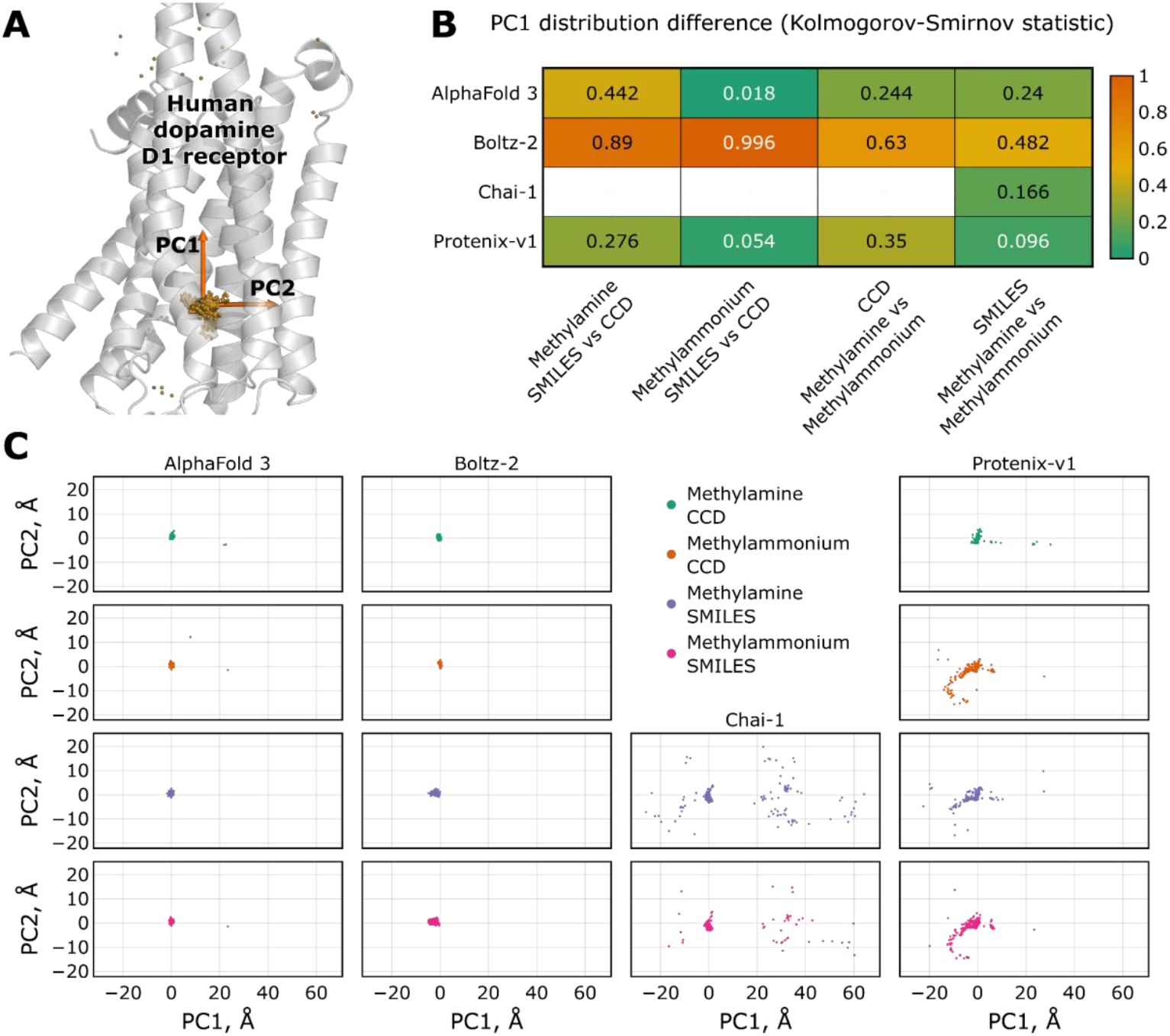
Predicted methylamine and methylammonium positions for the human dopamine D1 receptor. A) Overlay of the receptor structure (apo-form, predicted using AlphaFold 3) and predicted carbon atom positions (yellow). Predicted positions are predominantly observed around the expected amino group position in known ligands. PC1 and PC2 are the principal components for the carbon atom positions of the ligands. B) Values of the Kolmogorov-Smirnov statistic for pairs of carbon atom distributions projected on PC1, obtained using different tools. The metric ranges from 0 for identical distributions to 1 for non-overlapping ones. C) Carbon atom positions projected on PC1 and PC2 (A).

For the BarA sensor Cache domain, no experimental structure was publicly available at the moment of the analysis. Structural modeling, docking and mutagenesis place the ligand binding site in the vicinity of L95, T98, I130, T133 and I136.[29] None of the algorithms recapitulate this putative binding site (Fig. 4). For AlphaFold 3, predictions for acetate specified using CCD strongly differ from all other predictions; in all other cases the distributions are only slightly distinguishable.

**Figure 4.**
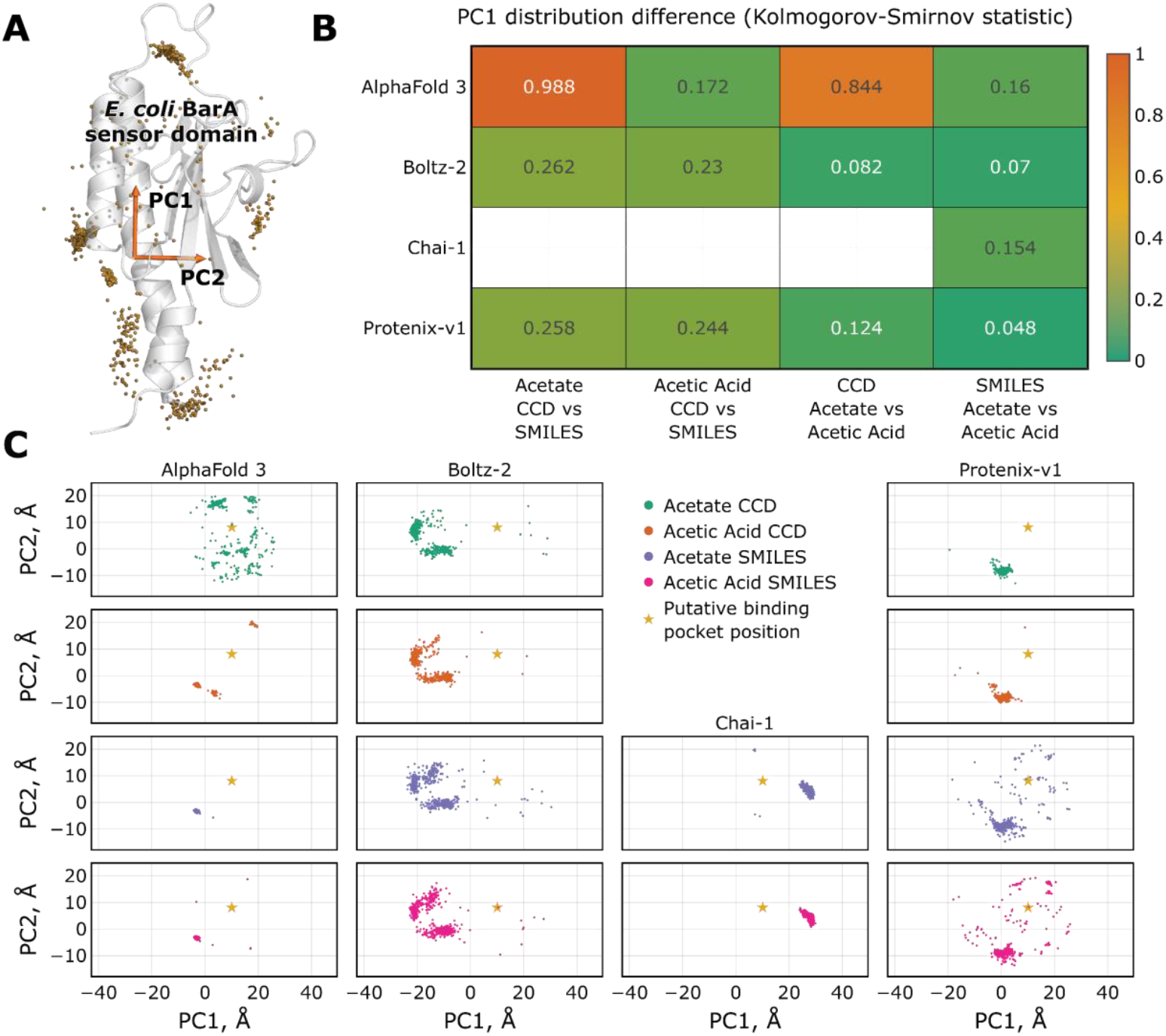
Predicted acetate and acetic acid positions for the *E. coli* BarA sensor domain. A) Overlay of the receptor structure (apo-form, predicted using AlphaFold 3) and predicted carboxyl carbon atom positions (yellow). PC1 and PC2 are the principal components for the carboxyl carbon atom positions of the ligands. B) Values of the Kolmogorov-Smirnov statistic for pairs of carboxyl carbon atom distributions projected on PC1, obtained using different tools. The metric ranges from 0 for identical distributions to 1 for non-overlapping ones. C) Carboxyl carbon atom positions projected on PC1 and PC2 (A). The expected position of the binding site [29] is marked with a star.

## Discussion

Advances in machine learning and artificial intelligence have led to development of numerous methods that excel at predicting folded protein structure, most prominently AlphaFold [2, 40]. These methods revolutionized structural biology [1] by providing high quality models across the protein universe [41] and highlighting the diversity of natural folds [42]. Current state-of-the-art approaches are also capable of producing models of protein-ligand complexes, which is useful both for studying natural proteins as well as for development of molecules targeting them [2, 3, 5, 6]. Yet, ligand binding site prediction by these tools is less reliable than protein structure prediction, becoming more complicated for chemical moieties underrepresented in training data [7, 12, 13].

One of the beneficial aspects of machine learning is that the resulting algorithm may sometimes generalize well and produce reasonable predictions for tasks it was not trained to do. For example, AlphaFold 2 was found to efficiently generate protein-protein complex models despite being initially trained on monomers [40, 43]. Accordingly, we decided to test whether the recently developed co-folding tools, AlphaFold 3, Boltz-2, Chai-1 and Protenix-v1, which use CCD and SMILES input formats, are able to produce structural models that take the charge of the ligand into consideration. We chose methylamine and acetic acid as two of the simplest titratable probes. These moieties are commonly found at the N- and C-termini of proteins, as well as in the side chains of lysine, aspartate and glutamate. As such, they are ubiquitous in the publicly available structures from the PDB [44], and undoubtedly present in the training sets of co-folding tools. Unexpectedly, we found that the results of the predictions generally depend stronger on the input format rather than on the formally specified charge of the ligand. On the other hand, charge dependence does not follow the expected pattern.

First of all, we found that the covalent bond lengths are on average shorter than expected from comprehensive quantum chemical calculations (Figs. 1 and 2, Table 1). The tools also differ in the spread of the values, with Protenix-v1 sometimes producing values as short as 0.075 Å; this may be fixed in future releases, as earlier Protenix v. 0.5.0 did not display such deviations. For protonated acetic acid, where the two C-O bonds are expected to be of different lengths due to different valency, none of the tools capture this effect (Fig. 2B). On the other hand, C-O bond lengths in acetate, where they are expected to be similar, are often systemically different from each other (Fig. 2B). Deviation of amino acid bond length and angle distributions from experimentally observed ones was previously described for AlphaFold 3 [10], so bond length deviations for ligands are not completely unexpected. Still, this is clearly not ideal and presents an opportunity for improving biomolecular structure prediction algorithms, especially given the availability of high quality molecular datasets and general purpose force fields.

Second, we tested the ability of the co-folding algorithms to place the small ligands at the expected binding sites. DRD1 binds numerous amines, with their amino group forming an ion bridge with the side chain of D103. Accordingly, we find that methylammonium is preferentially positioned at the expected position by all tools, with Chai-1 and Protenix-v1 predictions being slightly more diverse. For BarA, none of the co-folding algorithms recapitulate the expected configuration of the acetate binding site, which might be the consequence of absence of similar molecular arrangements in the training sets. Indeed, partially hydrophobic site binding a protonated carboxy group-containing ligand is not a common occurrence. Overall, our findings correspond to larger benchmarks that found a correlation between presence of a ligand in the training set and success rate of a corresponding prediction [7, 12, 13].

The next tested variable was the formally specified charge of the titratable ligands, correct treatment of which further complicates protein–ligand structure prediction [45]. Protonation cannot be deduced directly when using two of the most popular experimental structural biology methods, X-ray crystallography and cryo-electron microscopy, and thus the information on protonation of amino and carboxy groups is lacking from PDB. Hence, it is not unexpected that the algorithms trained on the data lacking the protonation information mix up different protonation forms of titratable amino acids. We find that even though the input formats allow direct specification of charge, resulting predictions do depend on charge, yet this dependence does not follow any pattern that would be expected from basic physical and chemical principles (Figs. 3 and 4). Ligand bond lengths do not change as expected from the change in protonation. Neutral methylamine is predicted to occupy roughly the same spots by all algorithms except Boltz-2; for Boltz-2, it is slightly displaced and reoriented compared to methylammonium. For Protenix-v1, it is predicted to occupy the expected position even more often than methylammonium, for which the position is more frequently shifted towards the extracellular surface of the protein. From general considerations, being essentially a hydrophobic compound, methylamine could be expected to be positioned in the hydrophobic regions of transmembrane receptor DRD1. For acetate, while the ligand was not predicted to bind at the putative site by any of the algorithms, only AlphaFold 3 produced different models for protonated and unprotonated forms when specified using CCD (Fig. 4C). Thus, correct accounting for the charge of amino and carboxy groups is also a clear area for improvement of the co-folding methods. One potential avenue is to apply physics-based refinement to the predicted ligand poses [46]. Alternatively, experimental structures can be augmented by assigning the protonation status and hydrogen positions using empirical rules, quantum mechanics/molecular mechanics calculations or machine learning [47], and respective algorithms can be retrained to learn to discern the different protonation forms. Potentially, this could also allow for correct modeling of dependence of structural properties on solution pH.

The last important question is that of the input format dependence (Figs. 1-4). It is definitely beneficial that different input formats are allowed in AlphaFold 3, Boltz-2 and Protenix-v1. However, it is completely unexpected that the prediction results are different when the ligand is specified using either CCD or SMILES. This discrepancy will probably be rectified in the next versions of the algorithms. Quite surprising is also that while the predictions do not reflect charge of the molecule as would be expected from physical and chemical principles, the results depend on the specified charge nonetheless. Overall, this raises an important point that if diverse predictions are desired while using the current versions of the methods, all possible input formats and molecular charges should be tested, whereas any conclusions derived from the charge-induced differences should be treated cautiously.

To summarize, none of the four popular cofolding methods that we compared clearly stood out in terms of prediction quality. Thus, the mentioned issues remain to be solved and present an opportunity for further improvement of biomolecular structure prediction approaches; in the meantime, co-folding results obtained using AlphaFold 3, Boltz-2, Chai-1 and Protenix-v1 should be interpreted with caution.

## Conclusions

Many of the physiologically relevant proteins and ligands can change their protonation state and, consequently, total charge. We find that while the popular co-folding methods allow the user to specify ligand protonation, the resulting predictions are not in accordance with basic physico-chemical principles. Moreover, chosen input format, CCD or SMILES, affects the results of the prediction significantly. This presents two clear paths for improvement of protein-ligand structure prediction algorithms: (i) ensuring identical results irrespective of the input formats, and (ii) inclusion of protonation-related steps into training and prediction pipelines. Eventually, this might allow for modeling of pH-related conformational and binding changes.

## Data availability

The datasets supporting the conclusions of this article (all generated models and associated data) are available in the Zenodo repository: doi.org/10.5281/zenodo.18664202.

## Author contributions

Conceptualization: A.B., P.O., I.G.

Investigation: A.B., P.O., I.G.

Supervision: P.O., I.G.

Visualization: A.B.

Writing – original draft: A.B., I.G.

Writing – review and editing: A.B., P.O., I.G.

## Acknowledgements

This work was supported by the Ministry of Science and Higher Education of the Russian Federation (agreement 075-03-2026-305, project FSMG-2025-0003). P.O. is a member of Guangdong Provincial Higher Education Institution Innovation Team Project (Grant #2025KCXTD055).

## Abbreviations

CCD: Chemical Component Dictionary
SMILES: Simplified Molecular Input Line Entry System
GPU: Graphics Processing Unit
JSON: JavaScript Object Notation
MSA: Multiple Sequence Alignment
PCA: Principal Component Analysis
DRD1: Dopamine D1 Receptor
PDB: Protein Data Bank

## References

1. Abriata LA (2024) The Nobel Prize in Chemistry: past, present, and future of AI in biology. Commun Biol 7:1409. 10.1038/s42003-024-07113-5

2. Abramson J, Adler J, Dunger J, et al (2024) Accurate structure prediction of biomolecular interactions with AlphaFold 3. Nature 630:493–500. 10.1038/s41586-024-07487-w

3. Discovery C, Boitreaud J, Dent J, et al (2024) Chai-1: Decoding the molecular interactions of life. 2024.10.10.615955

4. ByteDance AML AI4Science Team, Chen X, Zhang Y, et al (2025) Protenix - Advancing Structure Prediction Through a Comprehensive AlphaFold3 Reproduction. 2025.01.08.631967

5. Protenix Team, Zhang Y, Gong C, et al (2026) Protenix-v1: Toward High-Accuracy Open-Source Biomolecular Structure Prediction. 2026.02.05.703733

6. Passaro S, Corso G, Wohlwend J, et al (2025) Boltz-2: Towards Accurate and Efficient Binding Affinity Prediction. 2025.06.14.659707

7. Xu S, Feng Q, Qiao L, et al (2025) Benchmarking all-atom biomolecular structure prediction with FoldBench. Nat Commun 17:442. 10.1038/s41467-025-67127-3

8. Bret G, Sindt F, Rognan D (2026) Assessing Boltz-2 Performance for the Binding Classification of Docking Hits. J Chem Inf Model 66:1511–1521. 10.1021/acs.jcim.5c02630

9. Kim J, Correy GJ, Hall BW, et al (2025) Large scale prospective evaluation of co-folding across 557 Mac1-ligand complexes and three virtual screens. 2025.12.25.696505

10. Lyu N, Du S, Ma J, Herschlag D (2025) An Evaluation of Biomolecular Energetics Learned by AlphaFold. 2025.06.30.662466

11. Ishitani R, Moriwaki Y (2025) Improving Stereochemical Limitations in Protein–Ligand Complex Structure Prediction. ACS Omega 10:56075–56084. 10.1021/acsomega.5c07675

12. Menon KM, Davasam A, Chen G, et al (2025) AlphaFold3 for Structure-guided Ligand Discovery. 2025.12.04.692352

13. Škrinjar P, Eberhardt J, Tauriello G, et al (2025) Have protein-ligand cofolding methods moved beyond memorisation? 2025.02.03.636309

14. Masters MR, Mahmoud AH, Lill MA (2025) Investigating whether deep learning models for co-folding learn the physics of protein-ligand interactions. Nat Commun 16:8854. 10.1038/s41467-025-63947-5

15. Weininger D (1988) SMILES, a chemical language and information system. 1. Introduction to methodology and encoding rules. J Chem Inf Comput Sci 28:31–36. 10.1021/ci00057a005

16. Westbrook JD, Shao C, Feng Z, et al (2015) The chemical component dictionary: complete descriptions of constituent molecules in experimentally determined 3D macromolecules in the Protein Data Bank. Bioinformatics 31:1274–1278. 10.1093/bioinformatics/btu789

17. HMMER. http://hmmer.org/. Accessed 29 Jan 2026

18. Mirdita M, Schütze K, Moriwaki Y, et al (2022) ColabFold: making protein folding accessible to all. Nat Methods 19:679–682. 10.1038/s41592-022-01488-1

19. The UniProt Consortium (2025) UniProt: the Universal Protein Knowledgebase in 2025. Nucleic Acids Res 53:D609–D617. 10.1093/nar/gkae1010

20. Dimitropoulos D, Ionides J, Henrick K (2006) Using MSDchem to search the PDB ligand dictionary. Curr Protoc Bioinforma Chapter 14:14.3.1–14.3.21. 10.1002/0471250953.bi1403s15

21. Kim S, Chen J, Cheng T, et al (2025) PubChem 2025 update. Nucleic Acids Res 53:D1516–D1525. 10.1093/nar/gkae1059

22. Hamelryck T, Manderick B (2003) PDB file parser and structure class implemented in Python. Bioinformatics 19:2308–2310. 10.1093/bioinformatics/btg299

23. Pedregosa F, Varoquaux G, Gramfort A, et al (2011) Scikit-learn: Machine Learning in Python. J Mach Learn Res 12:2825–2830

24. Virtanen P, Gommers R, Oliphant TE, et al (2020) SciPy 1.0: fundamental algorithms for scientific computing in Python. Nat Methods 17:261–272. 10.1038/s41592-019-0686-2

25. The pandas development team (2026) pandas-dev/pandas: Pandas

26. Plotly (2024) Plotly PY

27. Schrödinger, LLC (2021) The PyMOL Molecular Graphics System, Version 2.5

28. Zhuang Y, Krumm B, Zhang H, et al (2021) Mechanism of dopamine binding and allosteric modulation of the human D1 dopamine receptor. Cell Res 31:593–596. 10.1038/s41422-021-00482-0

29. Alvarez AF, Rodríguez C, González-Chávez R, Georgellis D (2021) The Escherichia coli two-component signal sensor BarA binds protonated acetate via a conserved hydrophobic-binding pocket. J Biol Chem 297:101383. 10.1016/j.jbc.2021.101383

30. Chuev GN, Valiev M, Fedotova MV (2012) Integral Equation Theory of Molecular Solvation Coupled with Quantum Mechanical/Molecular Mechanics Method in NWChem Package. J Chem Theory Comput 8:1246–1254. 10.1021/ct2009297

31. Caskey DC, Damrauer R, McGoff D (2002) Computational studies of aliphatic amine basicity. J Org Chem 67:5098–5105. 10.1021/jo011118g

32. Chaiyasit P, Tongraar A, Kerdcharoen T (2017) Characteristics of methylammonium ion (CH3NH3+) in aqueous electrolyte solution: An ONIOM-XS MD simulation study. Chem Phys 493:91–101. 10.1016/j.chemphys.2017.06.012

33. Cabaleiro-Lago EM, Ríos MA (2000) Ab initio study of interactions in methylamine clusters. The significance of cooperative effects. J Chem Phys 112:2155–2163. 10.1063/1.480781

34. Gao D, Svoronos P, Wong PK, et al (2005) pKa of Acetate in Water: A Computational Study. J Phys Chem A 109:10776–10785. 10.1021/jp053996e

35. Rudolph WW, Fischer D, Irmer G (2014) Vibrational spectroscopic studies and DFT calculations on NaCH3CO2(aq) and CH3COOH(aq). Dalton Trans 43:3174–3185. 10.1039/C3DT52580E

36. Socha O, Dračínský M (2020) Dimerization of Acetic Acid in the Gas Phase—NMR Experiments and Quantum-Chemical Calculations. Molecules 25:2150. 10.3390/molecules25092150

37. Nakabayashi T, Kosugi K, Nishi N (1999) Liquid Structure of Acetic Acid Studied by Raman Spectroscopy and Ab Initio Molecular Orbital Calculations. J Phys Chem A 103:8595–8603. 10.1021/jp991501d

38. Abdi H, Williams LJ (2010) Principal component analysis. WIREs Comput Stat 2:433–459. 10.1002/wics.101

39. Berger VW, Zhou Y (2014) Kolmogorov–Smirnov Test: Overview. In: Wiley StatsRef: Statistics Reference Online. John Wiley & Sons, Ltd

40. Akdel M, Pires DEV, Pardo EP, et al (2022) A structural biology community assessment of AlphaFold2 applications. Nat Struct Mol Biol 29:1056–1067. 10.1038/s41594-022-00849-w

41. Varadi M, Anyango S, Deshpande M, et al (2022) AlphaFold Protein Structure Database: massively expanding the structural coverage of protein-sequence space with high-accuracy models. Nucleic Acids Res 50:D439–D444. 10.1093/nar/gkab1061

42. Lau AM, Bordin N, Kandathil SM, et al (2024) Exploring structural diversity across the protein universe with The Encyclopedia of Domains. Science 386:eadq4946. 10.1126/science.adq4946

43. Bryant P, Pozzati G, Elofsson A (2022) Improved prediction of protein-protein interactions using AlphaFold2. Nat Commun 13:1265. 10.1038/s41467-022-28865-w

44. Berman HM, Burley SK (2025) Protein Data Bank (PDB): Fifty-three years young and having a transformative impact on science and society. Q Rev Biophys 58:e9. 10.1017/S0033583525000034

45. Tosstorff A, Rudolph MG, Benz J, et al (2026) The CASP 16 Experimental Protein-Ligand Datasets. Proteins 94:79–85. 10.1002/prot.70053

46. Zhang L, Friesner RA, Miller EB, Rodrigues JP Generalization and Usability of Co-Folded GPCR–Ligand Complexes: A Physics-Guided Assessment. ChemRxiv 2026:. 10.26434/chemrxiv-2026-1rkqz

47. Wang Y, Sun K, Li J, et al (2025) A workflow to create a high-quality protein–ligand binding dataset for training, validation, and prediction tasks. Digit Discov 4:1209–1220. 10.1039/D4DD00357H

